# Single-cell analysis of lung epithelial cells reveals age and cell population-specific responses to SARS-CoV-2 infection in ciliated cells

**DOI:** 10.1101/2024.04.02.587663

**Authors:** Raven M. Osborn, Christopher S. Anderson, Justin R. Leach, ChinYi Chu, Stephen Dewhurst, Thomas J. Mariani, Juilee Thakar

## Abstract

The ability of SARS-CoV-2 to evade antiviral immune signaling in the airway contributes to the severity of COVID-19 disease. Additionally, COVID-19 is influenced by age and has more severe presentations in older individuals. This raises questions about innate immune signaling as a function of lung development and age. Therefore, we investigated the transcriptome of different cell populations of the airway epithelium using pediatric and adult lung tissue samples from the LungMAP Human Tissue Core Biorepository. Specifically, lung lobes were digested and cultured into a biomimetic model of the airway epithelium on an air-liquid interface. Cells were then infected with SARS-CoV-2 and subjected to single-cell RNA sequencing. Transcriptional profiling and differential expression analysis were carried out using Seurat.

The clustering analysis identified several cell populations: club cells, proliferating epithelial cells, multiciliated precursor cells, ionocytes, and two biologically distinct clusters of ciliated cells (FOXJ1^high^ and FOXJ1^low^). Interestingly, the two ciliated cell clusters showed different infection rates and enrichment of processes involved in ciliary biogenesis and function; we observed a cell-type-specific suppression of innate immunity in infected cells from the FOXJ1^low^ subset. We also identified a significant number of genes that were differentially expressed in lung cells derived from children as compared to adults, suggesting the differential pathogenesis of SARS-CoV-2 infection in children versus adults. We discuss how this work can be used to identify drug targets to modulate molecular signaling cascades that mediate an innate immune response and begin to understand differences in COVID-19 outcomes for pediatric vs. adult populations.

**Importance:** Viral innate immune evasion leads to uncontrolled viral spread in infected tissues and increased pathogenicity in COVID-19. Understanding the dynamic of the antiviral signaling in lung tissues may help us to understand which molecular signals lead to more severe disease in different populations, particularly considering the enhanced vulnerability of older populations. This study provides foundational insight into the age-related differences in innate immune responses to SARS-CoV-2, identifying distinct patterns of infection and molecular signaling in different cell populations of airway epithelial cells from pediatric and adult lung tissues. The findings provide a deeper understanding of age-related differences in COVID-19 pathology and pave the way for developing targeted therapies.

## Introduction

SARS-CoV-2, the virus responsible for the COVID-19 pandemic, is a positive-sense, single-stranded RNA virus that belongs to the coronaviridae family. SARS-CoV-2 binds host receptors for entry, and once inside the cell, the virus hijacks the host cell’s machinery to replicate its genome and produce viral proteins. Newly synthesized viral proteins and genomes assemble to create virions, which are then released from the cell where they can infect neighboring cells (1). During this process, host cell mechanisms for detecting the virus are in a race to prevent it from spreading to other cells. Viral infection induces secretion of type I and type III interferon, which leads to the induction of antiviral genes in the neighboring cells, making them resistant to viral infection through paracrine signaling (2–4). The balance between interferon secretion and viral spread determines the state of the innate immune response (2, 4).

A primary method for preventing the uncontrolled spread of SARS-CoV-2 after an organism has been infected is the robust induction of innate antiviral signaling cascades. Similar to other respiratory RNA viruses like Influenza, SARS-CoV-2 suppresses the antiviral host response. However, it has been demonstrated that SARS-CoV-2 can suppress the host’s antiviral response to a much greater extent than influenza in animal models (5). Furthermore, the antiviral interferon response has been inversely correlated with COVID-19 pathology (3, 6, 7).

Unsurprisingly, the inverse correlation between interferon secretion and severe coronavirus-related pathology is exacerbated in older age groups (7). In COVID-19, children generally experience milder acute illness; however, post-acute sequelae and Multisystem Inflammatory Syndrome in Children (MIS-C) occur in 1 of approximately 3,000 to 4,000 pediatric cases (8–10). One of the objectives of the present study was, therefore, to compare innate immune responses to SARS-CoV-2 in cells from children versus adults, with the goal of identifying and understanding age-related differences in these signaling pathways.

Interferon is the chief regulator of antiviral paracrine signaling, a mechanism by which cells adjacent to virus-infected cells can be driven into an antiviral state, thereby reducing viral spread. Many studies have interrogated this subject using cell lines, which don’t originate from the lung, or require the transduction of a known SARS-CoV-2 entry factor to support viral infection (5, 11, 12). Here, we have examined how these antiviral pathways are regulated after SARS-CoV-2 infection in primary human lung epithelial cells, using a biomimetic model cultured on an air-liquid interface that preserves the pseudostratification and differentiated functions of the normal lung epithelium.

SARS-CoV-2 infection is predominantly initiated and spread through cells in the airway epithelium. The respiratory epithelium comprises many different cell types, and SARS-CoV-2 virions infect those cells as they move throughout the respiratory system (13). Interestingly, the most commonly cited host entry factors for SARS-CoV-2 have heterogenous expression on cell-types along the respiratory system and within the cells of the same type (14). Mild and asymptomatic infections are typically restricted to ciliated and goblet cells in the nasal passages or upper respiratory tract (14), while more severe illnesses and pathology occur when the virus travels down to the parenchyma, infecting and injuring type II alveolar cells, disrupting the epithelial layer and reducing gas exchange – resulting in shortness of breath and respiratory distress (1). Therefore, investigating the host response to SARS-CoV-2 in cells of the conducting airway is important in understanding and preventing severe COVID-19 pathology. Due to the heterogeneity of cell types in the lung epithelium, we utilized single-cell RNA sequencing to discern cell-type-specific host responses in cultured primary lung epithelial cells.

## Materials and Methods

### Ethics statement

Donor lungs were provided through the federal United Network of Organ Sharing via the National Disease Research Interchange and the International Institute for Advancement of Medicine. With written consent, dissociated lung cells from deceased donors were entered into the LungMAP program’s biorepository and were utilized in this study. The University of Rochester’s Institutional Review Board approved and oversaw this study (RSRB00047606).

### Primary human cells

The LungMAP program’s biorepository was utilized in this study. The University of Rochester’s Institutional Review Board approved and oversaw this study (RSRB00047606). The study cohort included five infants (under six months old) and two adult donors (over fifty years old). All donors were male, two had an unknown race, and five were white. Pathologist notes from all donors showed normal lung structure; all pediatric donors had normal lung development, and both adult donors had some signs of chronic inflammation. None of the pathologist notes indicated a lung-associated cause of death.

### Viruses

The following reagents were deposited by the Centers for Disease Control and Prevention and obtained through BEI Resources, NIAID, NIH: SARS-Related Coronavirus 2, Isolate Hong Kong/VM20001061/2020, NR-52282. SARS-CoV-2 was propagated and titered using African green monkey kidney epithelial Vero E6 cells (American Type Culture Collection, CRL-1586) in Eagle’s Minimum Essential Medium (Lonza, 12-125Q) supplemented with 2% fetal bovine serum (FBS) (Atlanta Biologicals), 2 mM l-glutamine (Lonza, BE17-605E), and 1% penicillin (100 U/ml) and streptomycin (100 ug/ml). Virus stocks were stored at − 80°C. All work involving infectious SARS-CoV-2 was performed in the Biosafety Level 3 (BSL-3) core facility of the University of Rochester, with institutional biosafety committee (IBC) oversight.

### Cell culture on air-liquid interface

Primary human lung cells were cultured on an air-liquid interface as described (15, 16). Briefly, lung tissue issues were digested with a protease cocktail, and adherent cells were expanded with bronchial epithelial cell growth medium (Lonza, CC-3170) and then transferred to a collagen-coated transwell plate (Corning, 3470) until each well reached a transepithelial electrical resistance (TEER) measurement of >300 ohms. Cells were then placed on an air-liquid interface (ALI) by removing media from the apical layer of the transwell chamber and continuing to feed cells on the basolateral layer as they differentiate. Cells were differentiated for 4-5 weeks at ALI before experiment use.

### SARS-CoV-2 infections of airway epithelial cells

The apical layer of primary lung cells cultured at the air-liquid interface for 4-5 weeks was inoculated with SARS-CoV-2 (BEI, NR-52281, hCoV-19/USA-WA1/2020) at an MOI of 5 (titered in VeroE6 cells) in phosphate-buffered saline containing calcium and magnesium (PBS++; Gibco, 14040-133) and incubated at 37°C for 1.5 hours. Next, the infectious solution was removed, and the apical layer was washed with PBS++ (PBS with added calcium and magnesium). Cells were then incubated for 48 hours.

### SARS-CoV-2 inactivation and scRNA-seq sample preparation

Primary human lung cells infected with SARS-CoV-2 were prepared for scRNA-seq using a method described by this group (17, 18). Briefly, cultured cells were washed by dispensing and aspirating 37°C HEPES buffered saline solution (Lonza, CC-5022) and then dissociated with 0.025% Trypsin/EDTA (Lonza, CC-5012) for 10 min at 37°C. Dissociated cells were aspirated using a wide-bore pipette tip and placed in a tube containing ice-cold Trypsin Neutralization Solution (Lonza, CC-5002); this was repeated to maximize cell collection. Cells were then pelleted by centrifugation (300 x *g* for 5 min), resuspended in chilled HEPES, and centrifugally pelleted again. Next, the supernatant was removed using a wide-bore pipette tip, and the cell pellet was resuspended in 100 µl of chilled 1X DPBS. Next, 1 ml of a chilled 1:1 methanol acetone mixture was added to the cells dropwise with continuous gentle agitation. Cells were incubated on ice for 1 hour, washed in PBS++, counted, and finally resuspended in a cold SSC cocktail (3× Lonza AccuGENE SSC, BMA51205 + 0.04% BSA + 1mM DTT + 0.2 U/ µl RNase1 inhibitor).

### Library preparation and sequence mapping

Following SARS-CoV-2 inactivation and rehydration, cell suspensions were processed to generate single-cell RNA-Seq libraries using Chromium Next GEM Single Cell 3′ GEM, Library and Gel Bead Kit v3.1 (10x Genomics), per the manufacturer’s recommendations, as summarized below. To minimize the addition of the rehydration buffer, a maximum of 4 µl cell suspension was used in the GEM (Gel Bead-in-Emulsion) generation step. Subsequently, samples were loaded on a Chromium Single-Cell Instrument (10x Genomics, Pleasanton, CA, USA) to generate single-cell GEMs. GEM reverse transcription (GEM-RT) was performed to produce a barcoded, full-length cDNA from poly-adenylated mRNA. After incubation, GEMs were broken, the pooled GEM-RT reaction mixtures were recovered, and cDNA was purified with silane magnetic beads (DynaBeads MyOne Silane Beads, PN37002D, ThermoFisher Scientific). PCR further amplified the purified cDNA to generate sufficient material for library construction. Enzymatic fragmentation and size selection was used to optimize the cDNA amplicon size, and indexed sequencing libraries were constructed by end repair, A-tailing, adaptor ligation, and PCR. The final libraries contain the P5 and P7 priming sites used in Illumina bridge amplification. Sequence data were generated using Illumina’s NovaSeq 6000. Samples were demultiplexed and counted using 10x Cell Ranger version 6.0.1 with standard parameters. Samples were aligned against a combined reference containing the 10x provided human reference (GRCh38-2020-A) and NCBI GenBank SARS-CoV-2 reference sequence MT644268.

### Quality control and cell clustering

Analysis of scRNA-seq data was done using the Seurat v.4 R package (19). Cells expressing greater than 6000 genes or over 10% mitochondrial genes were omitted from the analysis. Sample integration was performed using the recommended scRNA-seq integration pipeline with canonical correlation analysis (CCA). Linear dimension reduction using principal component analysis (PCA) was then performed on the integrated data to determine the appropriate number of dimensions for Seurat’s clustering algorithm.

### Cell cluster annotation

Cell clusters were annotated using the Seurat v.4 R package (19). For each cluster, we performed the Wilcoxon Rank Sum test with a log fold change threshold of at least 0.25. Differentially expressed genes (DEGs) that are specific to each cluster were used for enrichment analysis with ToppGene Suite’s ToppCell Atlas to determine the cell type of each cluster (20). Functional analysis of DEGs was performed with the clusterProfiler 4.2 R package alongside Gene Ontology terms for biological processes (21). Cluster annotations were further performed for each cell using gene set enrichment analysis against a database of known cell type markers with the scType R package (22). Finally, cell assignment proportions were compared using the Chi-square test and visualized using the dittoSeq R package version 1.8.1 (23).

### Trajectory and pseudotime analysis

Normalized gene expression data, cluster assignments, UMAP embeddings, and partitions were converted from Seurat objects to a monocle object for trajectory and pseudotime analysis using the Monocle3 R package version 1.3.1 (24). Trajectory analysis was performed with the Seurat-assigned clusters on the entire dataset by assigning all cells to the same partition in Monocle3. Pseudotime was calculated using the basal cells as the root cluster when ordering cells. Pseudotime comparisons were performed with the Wilcoxon Rank Sum test with a Benjamini-Hochberg corrected adjusted p-value of less than .05.

### Cell-cell communication analysis

Cell-cell communication analysis was performed using the Liana framework from the Liana R package version 0.1.12 (25). A Robust Rank Aggregate score was calculated using algorithms from the following methods: NATMI, iTalk (logFC Mean), Connectome, SingleCellSignalR, and CellphoneDB (26–31). Context deconvolution was performed using the standard Liana framework in conjunction with Tensor-cell2cell and SingleCellSignalR (28, 32). Finally, footprint enrichment analysis with ligand-receptor pairs was performed using genesets from PROGENy and decoupleR R packages as recommended from the Liana context factorization pipeline (33, 34).

### Age and infection differential expression analysis

Differential gene expression analyses for age and infection were conducted using the MAST method in the FindMarkers function in the Seurat R package. The differentially expressed genes had log fold change >= 0.2 and a Benjamini-Hochberg corrected p-values <.05 (35). Differential pathway analysis was performed using the FindMarkers function using the MAST R package (version 1.22.0) with a Benjamini-Hochberg corrected p-value threshold of less than for the mock vs. infected contrast and .0001 for the age contrast (35). Cluster-specific intersections were visualized using the UpSetR (version 1.4.0) and ComplexUpset (version 1.3.3) R packages (36, 37).

### Gene functional annotation

Enrichment of DEGs from each cluster was performed and visualized using the clusterProfiler 4.2 R package (21). Over-representation analysis with clusterProfiler was used with genesets from the Kyoto Encyclopedia of Genes and Genomes (KEGG) 2021 pathway database with disease-associated pathway manually omitted (38). Gene set enrichment analysis was performed using the AUCell R Package version 1.18.1. AUCell scores were calculated using the top 10% of ranked genes in each cell against the KEGG genesets, a manually curated list of antiviral genes, and a geneset from a published overexpression screen of interferon-stimulated genes (39, 40). Cluster-specific intersections were visualized using the UpSetR (version 1.4.0) and ComplexUpset (version 1.3.3) R packages (36, 37).

## Results

### Primary lung epithelial cells cluster into six cell types, including two distinct ciliated cell clusters

Single-cell RNA sequencing (scRNA-seq) data from 52,482 cells from pediatric and adult donors with or without SARS-CoV-2 infection clustered into six cell populations (Fig. 1A) (Fig. 1A, SFig. 1). Annotation of these six cell populations using overrepresentation analysis (ORA) and gene set enrichment analysis (GSEA) revealed that primary human epithelial cells cultured on an air-liquid interface differentiated into FOXJ1^low^ ciliated cells, club cells, FOXJ1^high^ ciliated cells, basal cells, multiciliated precursor cells, and ionocytes (Fig. 1B-C). Functional enrichment analysis of cluster markers supported the underlying biological processes associated with those cell populations (Fig. 1D). Ciliated cell clusters had significant enrichment of pathways involved in cilium assembly, cilium organization, and microtubule-based movement (Fig. 1D). More transient cell populations (basal cells and multiciliated precursors) showed enrichment in pathways involving cell division (Fig. 1D). In total, our primary human epithelial cells cultured on an air-liquid interface differentiated into 35,279 (67.22%) FOXJ1^low^ ciliated cells, 9,194 (17.52%) club cells, 4,138 (7.89%) FOXJ1^high^ ciliated cells, 2,879 (5.49%) basal cells, 725 (1.38%) multiciliated precursor cells, and 267 (.51%) ionocytes (Fig. 1E-F). These proportions of cell populations were similar across most donors (Fig. 1F).

**Fig 1.**
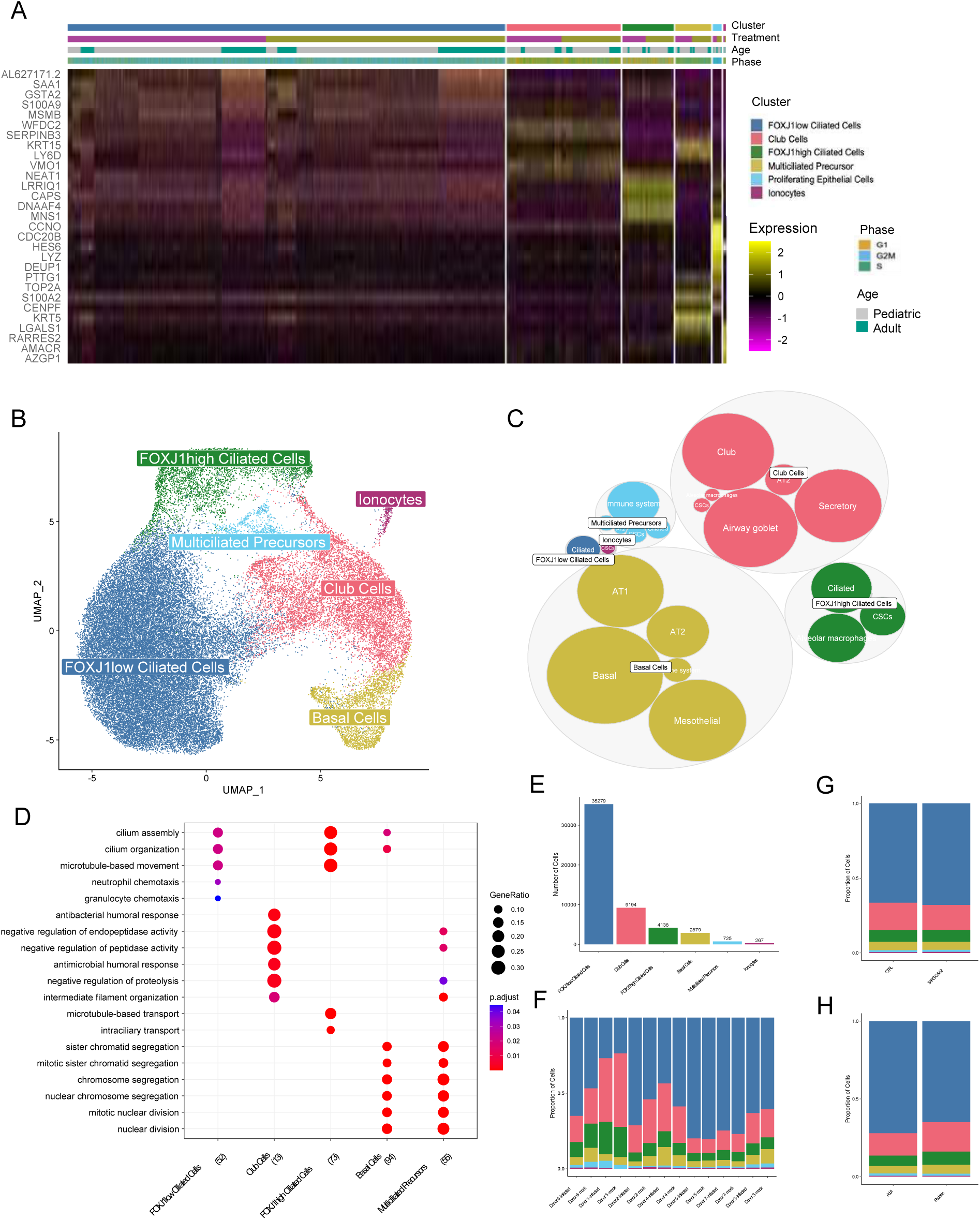
Clustering and annotation of scRNA-seq data from primary lung epithelial cells. (A) Heatmap of integrated gene expression of cluster markers from 52,482 cells annotated by assigned cluster, treatment status, donor age category, and cell cycle phase. (B) Uniform Manifold Approximation and Projection (UMAP) of all cells colored by cluster annotation. (C) Circular treemap of cell annotations. Circles are colored and clustered by higher-level annotations, and subclustered sizes are a function of the number of cells assigned to the subcluster. (D) Gene ontology (GO) analysis of cluster markers using Biological Process terms. (E) Bar plot of cell number in each cluster. (F) Stacked bar plot of cluster proportions for each sample. (G) Stacked bar plot of cluster proportions by treatment status. (H) Stacked bar plot of cluster proportions by age category.

We found that the proportion of cells in each cluster was the same in control samples as well as samples infected with SARS-CoV-2 (adjusted p>0.05)(Fig.1G)(SFig. 1A). Similarly, the proportions of cells in each population were not significantly different between pediatric and adult donors (Fig. 1H)(SFig. 1B). Further investigation of cell cycle scoring revealed that there were no significant differences in the proportion of cells in G1, G2M, or S phase between each donor, donor age, or treatment status (SFig. 2A-G). Although a high proportion of FOXJ1^low^ ciliated cells were found to have a G2M cell cycle score, this was due to low-level expression of a small subset of genes.

**Fig 2.**
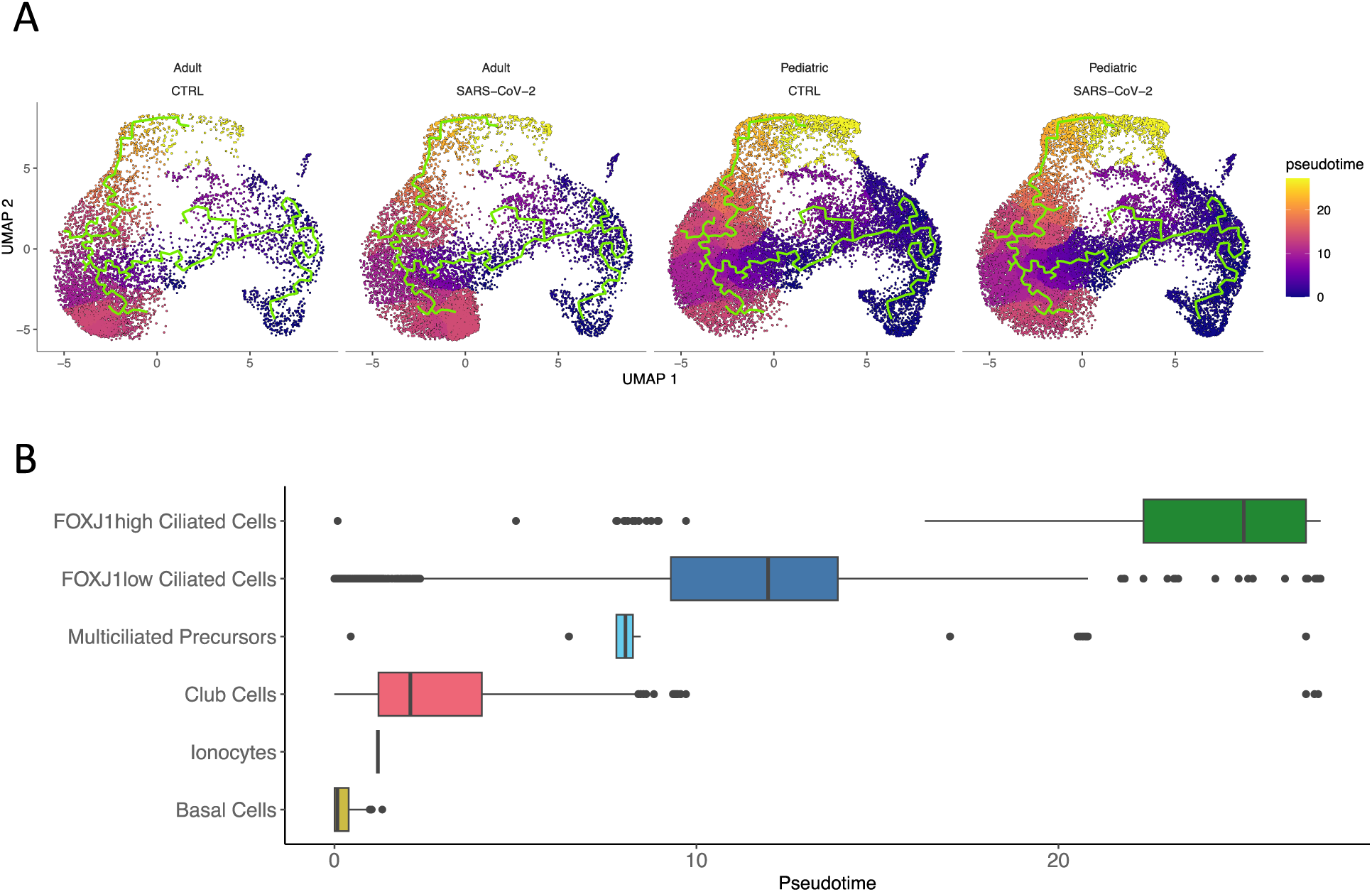
Trajectory analysis and pseudotime comparisons. (A) UMAPs separated by covariates with calculated trajectory in chartreuse, and colored by pseudotime representing differentiation from dark blue to yellow (more differentiated). (B) Boxplots of the range of pseudotime in each cluster, colored by cluster annotation shown in Figure 1.

The lung epithelium primarily consists of fully differentiated cells, but injury can cause cells to dedifferentiate into their progenitors (41–43). We were interested to see if there were differences in cell differentiation among age cohorts or treatment status. Trajectory analysis of the main partition revealed that the differentiation trajectory of our cells began with the most “stem-like” cell type, basal cells, and continued onto club cells, multiciliated precursors, FOXJ1^low^ ciliated cells, and finally FOXJ1^high^ ciliated cells (Fig.2). Trajectories of all the cells from each sample showed no obvious differences in cell trajectories between any age or treatment status. Further, we found that the two ciliated cell types were related and did not arise from two different branches. Rather, FOXJ1^high^ ciliated cells arise from FOXJ1^low^ ciliated cells. Indeed, our pseudotime analysis revealed that FOXJ^high^ ciliated cells were the most terminally differentiated cell type, followed by FOXJ1^low^ ciliated cells, multiciliated precursors, club cells, ionocytes, and finally basal cells (which were defined as the root cells in our analysis) (42–44) (Fig. 2)(SFig. 3).

**Fig 3.**
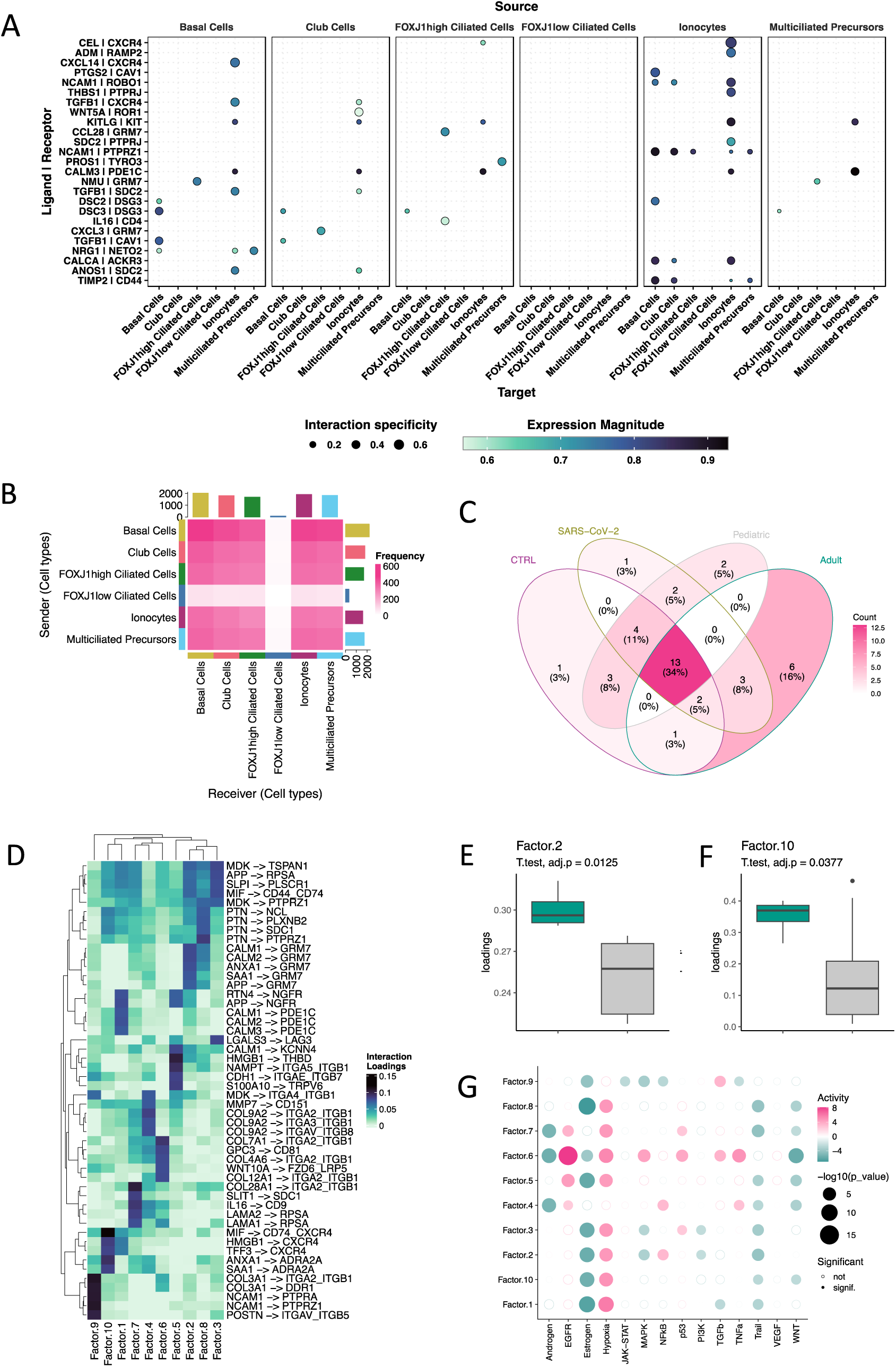
Cell-cell communication analysis. (A) Dot plots of ligand-receptor pairs (y-axis) show the source cell type sending the signal on top and the receiving cell populations on the bottom. (B) Heatmap of the frequencies of interactions for each pair of potentially communicating cell types. Annotation bar plot on top (“receiving”) and right (“sending”) is the total number of interactions per cell type. (C) Venn diagram of top 25 ligand-receptor pairs across infection and age. (D) Heatmap of top 5 ligand-receptor loadings for each Factor. (E and F) Boxplots of age-related factor loadings with adult samples in green and pediatric samples in grey. (G) Dotplot of footprint enrichment analysis of downstream pathways.

### FOXJ1^low^ ciliated cells exhibit distinctly low ligand-receptor communication patterns

Intercellular communication is vital in preserving homeostasis and serves as one of the primary mechanisms for eliciting an immune response in complex biological systems such as the airway epithelium. To investigate intercellular communication in our primary cells, we aggregated cell-cell communication scores from several methods. Ionocytes had many inferred ligand-receptor interactions when top receptors are ordered by magnitude and then specificity; these interactions occur with ionocytes, suggesting regulatory feedback, or basal cells (Fig 3A).

When quantifying the frequency of inferred outgoing signals through ligand expression, basal cells had the highest frequency (2201), followed by club cells (1809), multiciliated precursors (1739), FOXJ1^high^ ciliated cells (1695), ionocytes (1601), and FOXJ1^low^ ciliated cells (339)(Fig. 3B)(Table 1). Basal cells were also found to have the highest frequency of inferred signal reception (2016) followed by ionocytes (1917), multiciliated precursors (1843), club cells (1815), FOXJ1^high^ ciliated cells (1698), and FOXJ1^low^ ciliated cells (95)(Fig. 3B)(Table 1)(SFig. 4). Interestingly, the most numerous cell population, FOXJ1^low^ ciliated cells, had very few inferred intrapopulation and interpopulation ligand-receptor interactions as a sender or receiver (Fig. 3B)(Table 1). However, FOXJ1^low^ ciliated cells were ∼3.5 times more likely to send signals rather than receive them (Fig. 3B)(Table 1).

**Fig 4.**
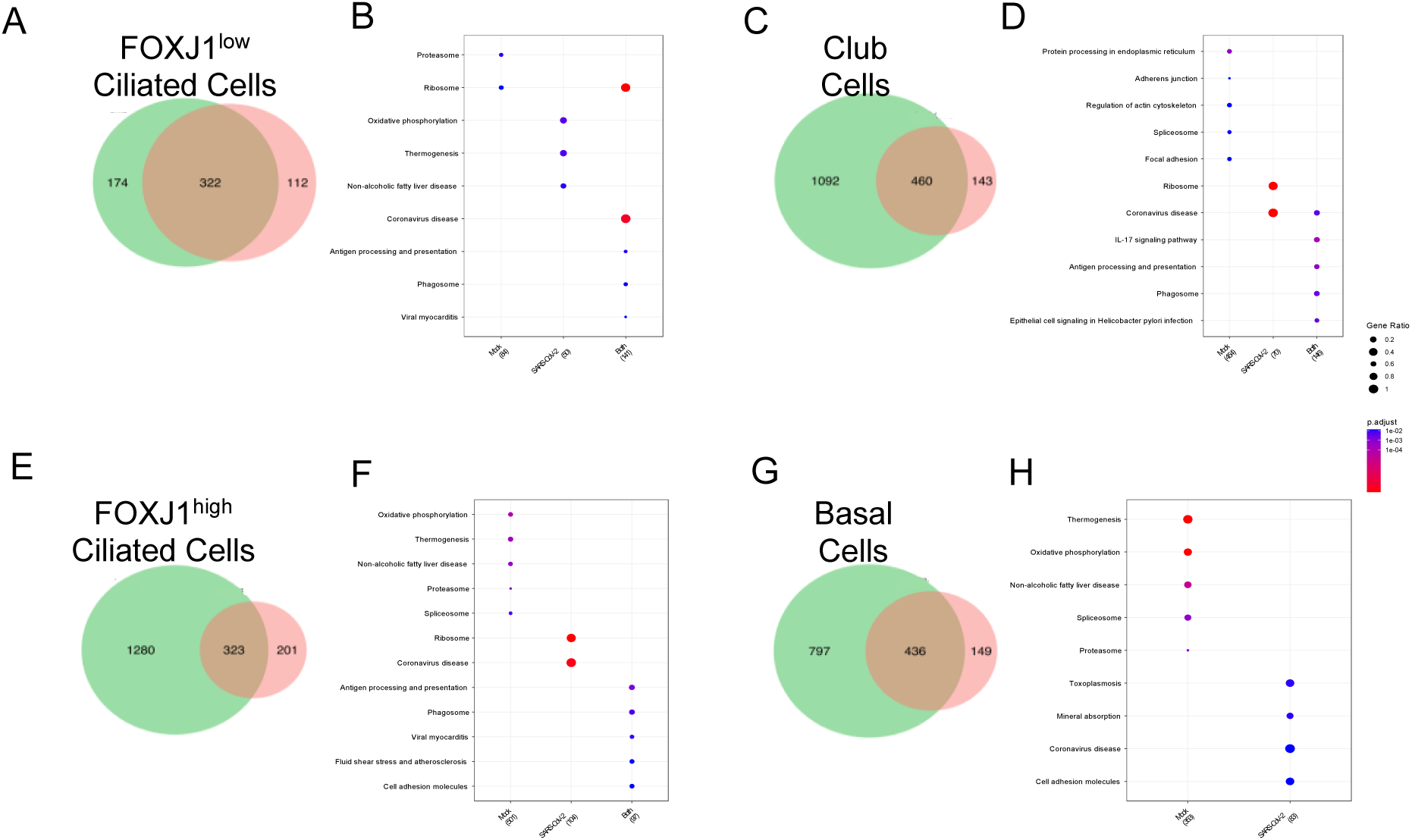
Age-related differential expression analysis. (A-H) Venn diagrams and dot plots of differentially expressed genes in pediatric vs adult donors present in mock (CTRL, green) and SARS-CoV-2-infected (red) samples for FOXJ1^low^ ciliated cells (A and B), club cells (C and D), FOXJ1^high^ ciliated cells (E and F), and basal cells (G and H).

**Table 1.** Sender and receiver cell frequencies. Frequency table of the predicted sender and receiver cell communication.

|  | Combined<br>Senders | Combined<br>Receivers | CTRL<br>Senders | CTRL<br>Receivers | SARS-CoV-2<br>Senders | SARS-CoV-2<br>Receivers | Pediatric<br>Senders | Pediatric<br>Receivers | Adult<br>Senders | Adult<br>Receivers |
| --- | --- | --- | --- | --- | --- | --- | --- | --- | --- | --- |
| FOXJ1low Ciliated<br>Cells | 339 | 95 | 362 | 94 | 333 | 90 | 311 | 89 | 433 | 119 |
| Club Cells | 1809 | 1815 | 1881 | 1916 | 1757 | 1778 | 1627 | 1746 | 2104 | 2217 |
| FOXJ1high Ciliated<br>Cells | 1695 | 1698 | 1661 | 1853 | 1653 | 1676 | 1623 | 1704 | 1821 | 1838 |
| Basal Cells | 2201 | 2016 | 2247 | 2081 | 2145 | 1981 | 2098 | 1910 | 2385 | 2175 |
| Multiciliated<br>Precursors | 1739 | 1843 | 1890 | 1868 | 1689 | 1734 | 1730 | 1638 | 1868 | 2022 |
| Ionocytes | 1601 | 1917 | 1812 | 2041 | 1550 | 1868 | 1529 | 1831 | 1709 | 1949 |

When comparing the top 25 most significant ligand-receptor pairs identified by all the cell-cell communication algorithms, we found 13 (34%) pairs common across age and infection comparisons. Additionally, there were few context-specific ligand-receptor pairs for each group; one in mock samples (C3-GRM7), one in SARS-CoV-2-infected samples (PROS1-TYRO3), two in pediatric samples (GDF15-RET, VEGFB-RET), and six in adult samples (MMP1-CD44, TGFB1-CAV1, TGFB1-SDC2, TIMP3-CD44, DSC3-DSG3, SDC2-PTPRJ)(Fig. 3C).

To better understand the drivers behind these age or treatment-dependent differences, we performed cell-cell communication analysis on each sample to generate a tensor of ligand-receptor interactions and decomposed the tensor into communication patterns. The ionocyte cluster did not have enough representation in all samples, so we excluded ionocytes from samples with low counts in downstream analysis. After tensor deconvolution, we found ten significant factors that inform cell-cell communication patterns with distinct ligand-receptor pair expression patterns (Fig. 3D)(SFig. 5). Factor loadings were compared between treatment status and age. Surprisingly, we found no significant differences between treatment statuses. Still, factor 2 and factor 10 were significantly different between age groups (Fig. 3E)(Fig. 3F)(SFig. 6)(SFig. 7). Further exploration of the pathways associated with each factor revealed that factor 2 and 10 were downstream signaling typically involved in an antiviral response (Fig. 3G). Factor 2 was associated with increased NFκB activity and low MAPK, PI3K, and TRAIL signaling, while factor 10 was associated with reduced TRAIL signaling. These age-related differences in antiviral signaling prompted us to explore the differences in our age cohorts.

**Fig 5.**
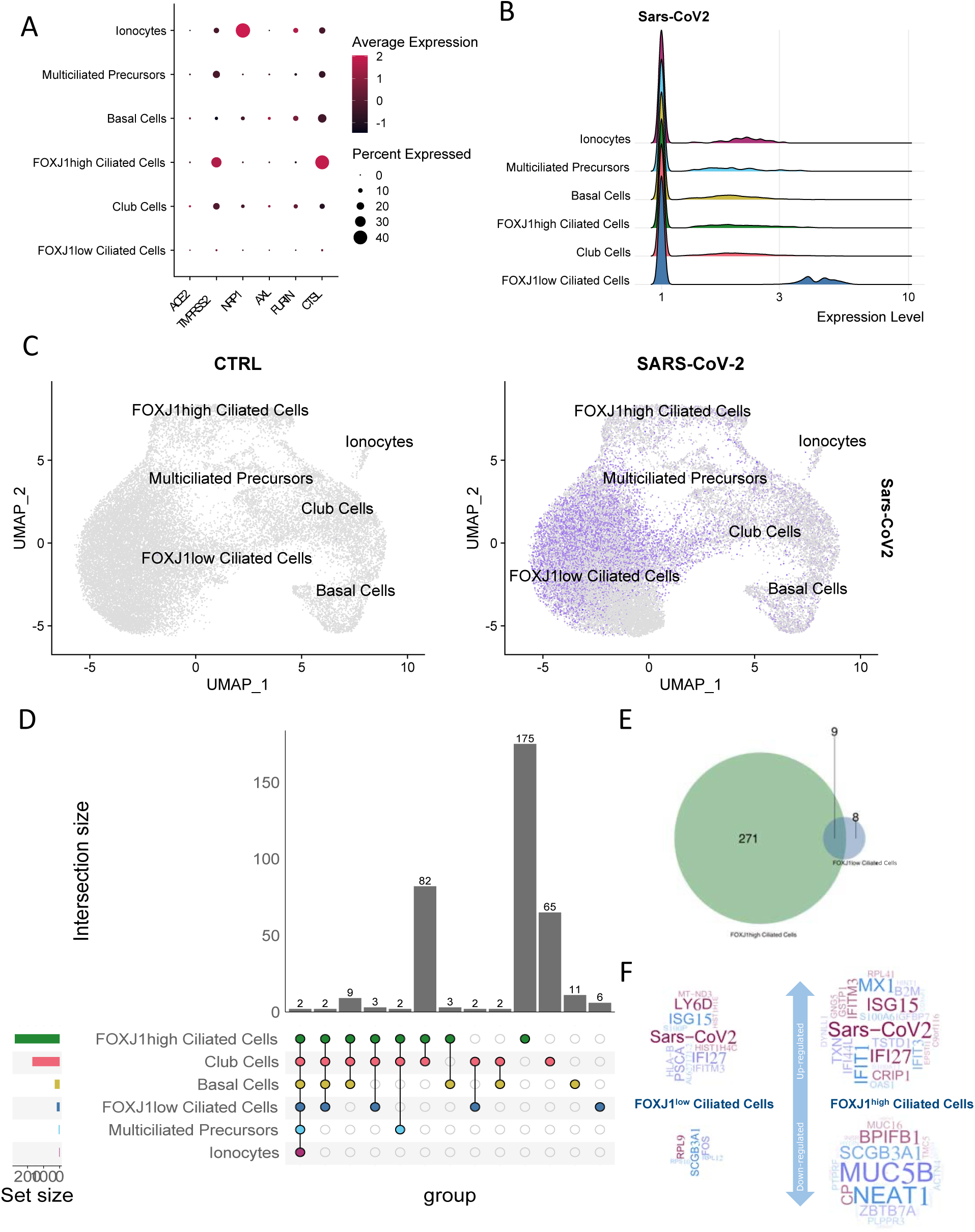
Cell-type-specific responses to SARS-CoV-2 infection. (A) Dotplot of scaled average gene expression for indicated SARS-CoV-2 host entry factors. (B) Ridge plot of log2 normalized SARS-CoV-2 expression in each cluster. (C) UMAPs of log2 normalized SARS-CoV-2 expression (purple) in each cell, split by treatment status. (D) An upset plot of DEGs in mock vs. infected cells by cluster. (E) Venn diagram of the intersection of treatment-related DEGs between ciliated cell clusters. (F) Word clouds of DEGs in ciliated cell clusters. The font size of the gene names is a function of the log fold change between CTRL and SARS-CoV-2 samples.

**Fig 6.**
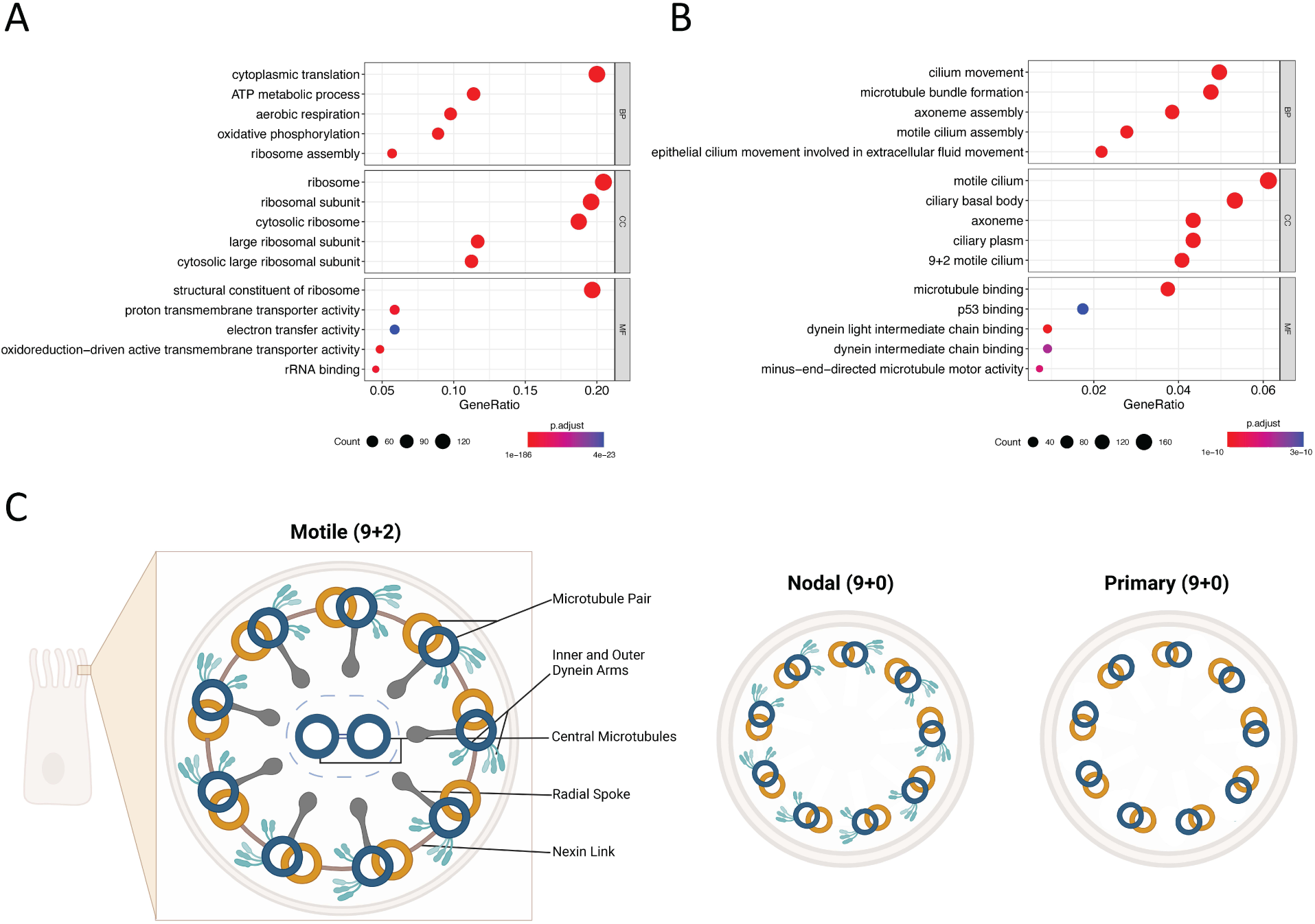
Ciliated cell functional comparison. (A and B) Dot plots of differentially enriched GO terms in FOXJ1^low^ ciliated cells compared to FOXJ1^high^ ciliated cells (A) and enriched terms in FOXJ1^high^ ciliated cells compared to FOXJ1^low^ ciliated cells (B). (C) Diagram of a cross-section of a ciliary axoneme. Three ciliary arrangements are shown: motile cilium (9+2) on the left, nodal cilium (9+0) in the center, and primary cilium (9+0) on the right.

**Fig 7.**
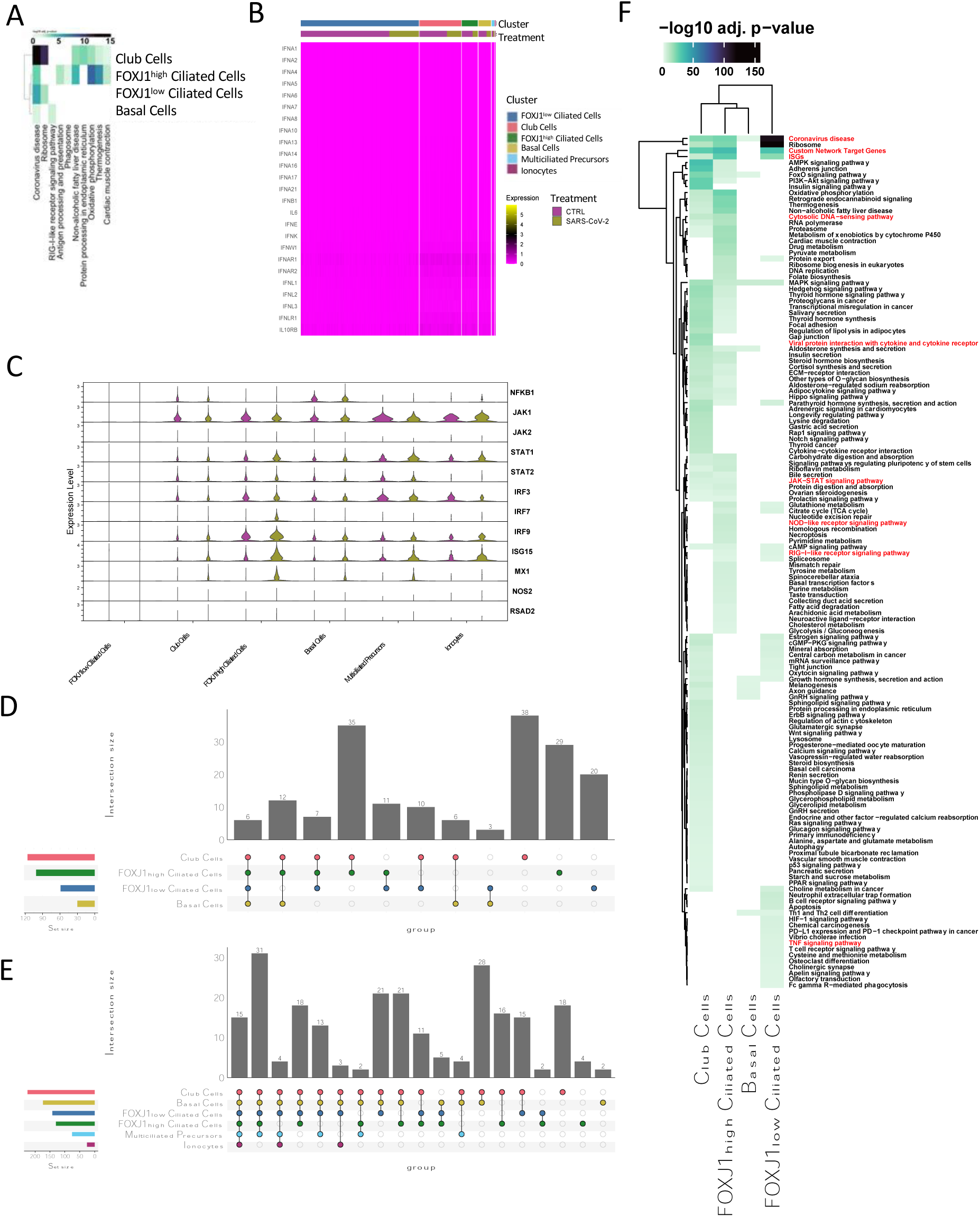
SARS-CoV-2 infection and cell-type-specific host responses. (A) Heatmap of -log10 adjusted p-values of differentially enriched KEGG pathways by cluster (y-axis) between mock and SARS-CoV-2-infected samples. (B) Heatmap of log2 normalized expression of indicated genes associated with a type I and type III interferon response, annotated by cluster and treatment status. (C) Violin plot of major transcription factors and intermediary genes associated with type I and III interferon responses. (D and E) An upset plot of differentially enriched AUCell scores of KEGG pathways between mock and SARS-CoV-2 treated samples (D) and pediatric vs. adult samples (E) by cluster. (F) Heatmap of -log10 adjusted p-values of significant differentially enriched AUCell scores of KEGG pathways by cluster between mock and SARS-CoV-2-infected cells. Pathways in red are typically associated with antiviral responses.

### Age-related differences varied by cluster and infection status

Comparisons of cluster-specific gene-expression changes revealed several differences between adult and pediatric donors. Using only the cells from mock-infected samples, we found that the total number of DEGs and the number of unique DEGs varied between clusters; notably, only a single gene was detected across all 6 cell types (RPS26), and only 22 genes were detected when we excluded the smallest cluster, ionocytes, from this analysis (SFig. 7A). Similar to the uninfected samples, we found that the total number of DEGs and the number of unique DEGs varied between clusters. There were 81 genes that were unique to ciliated cell populations, while basal and club cells had the highest combined number of DEGs in common. There were only three genes (RPS26, AL627171.2, and SARS-CoV-2) differentially expressed between adult and pediatric donors across all cell populations (SFig. 7B). Geneset enrichment analysis of combined mock and infected data showed that age-related DEGs in ciliated cells were highly related to ribosomal activity. In contrast, ciliated cells, basal cells, club cells, and multiciliated precursor cells showed differences in the coronavirus disease and antigen processing and presentation genesets (SFig. 8D)(SFig. 8E).

When comparing the age-related DEGs for each cluster in infected and mock samples, we found some interesting functional differences. The majority of age-related DEGs in FOXJ1^low^ ciliated cells were found in both mock and infected samples, while all other cell populations showed a stronger age-related difference in mock samples (Fig. 4A, C, E, and G). In FOXJ1^low^ ciliated cells, age-related DEGs had the strongest association in “Coronavirus disease” and “Ribosome” pathways, which were enriched in both treatment conditions alongside the “Antigen processing and presentation” pathway (Fig. 4B). Club cells, however, showed an age-related enrichment in “Coronavirus disease” and “Antigen processing and presentation” in both treatments, but “Ribosome” pathway enrichment was only found in the infected samples. On the other hand, we observed an age-related enrichment in the “Ribosome” and “Coronavirus disease” pathways in infected samples, while “Antigen processing and presentation” was enriched in both treatments from FOXJ1^high^ ciliated cells. Conversely, basal cells had little overlap in pathway enrichment between mock and infected samples (Fig. 4G, H)

Broadly, the large numbers of DEGs between adult and pediatric populations narrowed in SARS-CoV-2-infected samples compared to mock samples (Fig. 4A, C, E, and G). Cluster-specific functional analysis of age-associated DEGs in mock only, infected only, and in both sample sets showed that many of the age-related DEGs in control samples were related to protein synthesis, while age-related DEGs in infected samples were metabolism-related (Fig. 4B, D, F, and H). In addition, the largest clusters (FOXJ1^low^ ciliated cells, club cells, and FOXJ1^high^ ciliated cells) all had age-related differences in “Antigen Processing and Presentation” in mock and infected samples. Perhaps the most striking finding to emerge from these analyses was that the annotated transcript for SARS-CoV-2 was consistently higher in pediatric samples compared to adult samples, with the highest difference between pediatric vs. adult samples being detected in FOXJ1^low^ ciliated cells (a 3.120 log fold change).

### FOXJ1^low^ ciliated cells are highly infected by SARS-CoV-2

Common SARS-CoV-2 entry factors ACE2, TMPRSS2, NRP1, AXL, FURIN, and CTSL showed heterogeneous expression levels across cell populations. The commonly cited entry factor, ACE2, had very low expression in all clusters. Interestingly, TMPRSS2 and CTSL were the most commonly expressed entry factors, with around 10-30% of cells expressing them in all clusters with the highest distribution and average expression level in FOXJ^high^ ciliated cells (Fig. 5A). When comparing the expression level and percentage of cells expressing each gene by infection status in each cluster, there were no significant differences in gene expression among common SARS-CoV-2 entry factors ACE2, TMPRSS2, NRP1, AXL, FURIN, CTSL (SFig. 9A). Despite evidence of low expression of SARS-CoV-2 entry factors in cells from both mock and infected samples, FOXJ1^low^ ciliated cells had the highest overall expression of SARS-CoV-2 transcripts in qualitative and quantitative analysis (Fig. 5B)(Fig. 5C).

Comparisons of mock and infected samples resulted in the identification of cluster-specific infection-related DEGs. Similar to our age-related analysis, we found that the total number of DEGs and the number of unique DEGs varied between clusters; FOXJ1^low^ ciliated cells had 17 total and 6 unique genes (.35 distinct proportion), club cells had 170 total and 65 unique genes (.38 distinct proportion), FOXJ1^high^ ciliated cells showed 280 total and 175 unique genes (.63 distinct proportion), basal cells had 30 total and 11 unique genes (.37 distinct proportion), multiciliated precursor cells had 5 total and 0 unique genes, while ionocytes showed 2 total and 0 unique genes (Fig. 5D). We found only two genes that were differentially expressed between mock and infected samples across all cell populations; the transcript for SARS-CoV-2 and SCGB3A1, a gene that encodes a secreted protein involved in regulating epithelial cell proliferation, differentiation, morphogenesis, and defines secretory cell subsets (45, 46) (Fig. 5D). Strikingly, there were no ciliated-cell-specific DEGs between mock and infected samples. Moreover, ciliated cells only had 9 infection-related DEGs in common (Fig. 5E).

Investigation into the DEGs revealed that FOXJ1^low^ ciliated cells (the most susceptible to SARS-CoV-2 infection) had a muted antiviral response to SARS-CoV-2. In contrast, FOXJ1^high^ ciliated cells (which were less susceptible to the virus) had DEGs associated with a robust antiviral response (ISG15, MX1, and IFI27), dysregulated mucus secretion (MUC5B and BPIFB1), and down-regulated ciliogenesis (HYDIN) (Fig. 5F).

Stark differences in the host responses from each ciliated cell type prompted us to perform differential functional analysis between the two ciliated cell types. Metabolic and translation-relevant signaling pathways were significantly different between FOXJ1^low^ ciliated cells and FOXJ1^high^ ciliated cells (Fig. 6A). However, FOXJ1^high^ ciliated cells had more significant enrichment of pathways involved in ciliogenesis and cilia motility (Fig. 6B). These differences in biological functions are typically indicated by cilium subtypes, which is evident in the microanatomy of the cilium itself (Fig. 6C). FOXJ1^high^ ciliated cells had enrichment in axoneme assembly, dynein binding, and microtubule motor movement, typical of 9+2 cilium microanatomy (Fig. 6B). In contrast, FOXJ1^low^ ciliated cells were not enriched in the compartments and showed a much lower level of canonical ciliated cell marker expression (Fig. 6B)(SFig. 10A) (SFig. 10B).

### Cell-type-specific responses to SARS-CoV-2 infection

After finding many differences in the DEGs, we were interested in understanding how cluster-specific DEGs contributed to functional responses. Functional enrichment of each cluster’s DEGs after infection showed a stark difference in enriched pathways (Fig. 7A). Despite FOXJ1^low^ ciliated cells having the highest proportion of infected cells, all cell populations showed significant enrichment of the “Coronavirus disease” pathway, with the lowest p-value in club cells. Overall, FOXJ1^high^ ciliated cells had a response pattern more similar to club cells than FOXJ1^low^ ciliated cells (Fig. 7A). When interrogating the expression of interferon-related transcription factors and key genes, we found that all cell populations had low expression levels of factors mediating an interferon response regardless of infection status. (Fig. 7B). Also, no clusters significantly differed in type 1 or 3 interferon-related genes in infected samples compared to mock (Fig. 7C).

Further, we extended the pathway analysis by incorporating all expressed genes for each cell independently (see methods). The comparison of the cell-specific pathway activities found more differences across age than the infection status (Fig. 7D and E). The ciliated cell clusters, club cells, and basal cells had significantly different pathway activities across infection status, with club cells having the most (38 pathways); we detected only 6 common pathways between all clusters, including the “Coronavirus disease” pathway (Fig. 7D). Both ciliated populations and club cells had significant activity in “ISGs,” a previously described list of over 200 interferon-stimulated genes (Fig. 7F)(39, 40). Pathway scoring at the single cell level revealed that FOXJ1^low^ and FOXJ1^high^ ciliated cells both had significant activity in the “Rig-I-like receptor signaling pathway” along with basal cells (Fig. 7A and F). Club cells and FOXJ1^high^ ciliated cells had significant activity in “Cytosolic DNA-sensing” and “JAK-STAT signaling” pathways, while FOXJ1^high^ ciliated cells were the only population to have activity in the “NOD-like receptor signaling pathway”(Fig. 7F). FOXJ1^low^ ciliated cells were the only population to have significant activity in “TNF signaling” (Fig. 7F). There were 15 shared age-associated differences in pathways all clusters, and an additional 31 shared pathways between all clusters excluding the smaller ionocyte population (Fig. 7E). Club and basal cells had the most differences between age cohorts, and shared the most differentially active pathways (Fig. 7E). There were dramatic differences in the “ISGs” and “Antigen processing and presentation” pathways in all major cell populations (both ciliated cell clusters, club cells, and basal cells), “Coronavirus disease” in club cells and FOXJ1^low^ ciliated cells, and “Type 1 IFNs” in club cells only (SFig. 10).

## Discussion

Understanding the host response to SARS-CoV-2 infection is critical in developing effective strategies for preventing and treating COVID-19. The lung epithelium is the primary target of SARS-CoV-2 infection, and early prevention strategies targeting these cells can therefore be expected to determine disease severity and outcome. Inefficient or dysregulated host responses to SARS-CoV-2 infection in lung epithelial cells can lead to severe lung damage and respiratory failure, which are the leading causes of mortality in COVID-19 patients. Moreover, some of the earliest epidemiological evidence from the COVID-19 pandemic showed that age is a significant risk factor for severe COVID-19 (47–50). Therefore, investigating age-related host responses in lung epithelial cells is crucial for understanding the underlying mechanisms of increased susceptibility, pathogenesis, and severity of COVID-19 in different populations. Here, we employed single-cell RNA sequencing (scRNA-seq) to investigate the transcriptomic profiles of primary lung epithelial cells from pediatric and adult populations in response to SARS-CoV-2. Our scRNA-seq analysis revealed six cell types in air-liquid interface cultures derived from primary human lung tissue: two ciliated cell types (FOXJ1^low^ and FOXJ1^high^), club cells, basal cells, multiciliated precursors, and ionocytes (Fig. 1).

Trajectory and pseudotime analysis were used to investigate the differentiation pathways of various cell types in our cell culture system. Basal cells are known as the most “stem-like” cells in the lung epithelium (43, 44). Trajectory analysis showed that basal cells differentiated into all of the other cell populations and confirmed that the ciliated cell populations were not two distinct branches of ciliated cells; instead, they were at different levels of differentiation, with FOXJ1^high^ ciliated cells being the most terminally differentiated. Density plots of 7 canonical ciliated cell markers confirmed that the most differentiated ciliated cells were located within the FOXJ1^high^ ciliated cell cluster (SFig. 10).

One drawback of using scRNA-seq analysis to understand viral infections on adherent cells is the absence of spatial data. This is especially true as infection-induced paracrine signaling elicits responses in the neighboring cells. Cell-cell communication analysis attempts to recapitulate multicellular coordination by inferring intercellular communication from the expression of genes associated with such communication (25). Using aggregated scores from various cell-cell communication methods, we found that FOXJ1^low^ ciliated cells had very little interpopulation and intrapopulation communication. While we expected cell communication patterns or expression of specific ligand-receptor pairs to change between infected and uninfected samples, no significant differences were observed. Notably, age was a significant factor in communication patterns (Fig. 3E)(Fig. 3F). Overall, footprint analysis of the ligand-receptor pairs in the factors that were different between age cohorts revealed that adult donors were more associated with cell communication patterns that resulted in signaling leading to high NFκB activity, and low MAPK, PI3K, and TRAIL activity in the adult donors (Fig. 3G). To our knowledge, this finding is novel in human age-related pulmonary biology.

Our data reveals cluster-specific differences in adult and pediatric samples (SFig. 8). When splitting the groups by infection status to remove infection-related variability, we were surprised to find that age-related differences were more prevalent in uninfected samples compared to infected samples in every cluster (Fig. 4A)(Fig. 4B)(Fig.S10). We found that the geneset for “Coronavirus disease” was differentially enriched between the age cohorts in both mock and infected samples in FOXJ1^low^ ciliated cells and club cells, while FOXJ1^high^ ciliated and basal cells only had age-related enrichment in the infected samples (Fig. 4B)(Fig. 4D)(Fig. 4F)(Fig. 4H). Generally, we interpreted this age-related

Similar to recent publications assessing the expression of SARS-CoV-2 entry factors along the airway epithelium, we found that our cells had very low expression of ACE2, TMPRSS2, NRP1, AXL, FURIN, and CTSL entry factor genes (Fig. 5A)(14, 51). Although this low expression was sufficient for SARS-CoV-2 infection in all cell populations, with the majority of infected cells belonging to the FOXJ1^low^ ciliated cell population (Fig. 5B)(Fig. 5D). However, FOXJ1^high^ ciliated cells showed the most robust responses to infection and followed more canonical antiviral signaling responses (Fig. 5D)(Fig. 7A) (Fig. 7F). Overall, FOXJ1^low^ ciliated cells seem to have inhibited transcription at the global level, leading to little overlap between the DEGs after infection between the two ciliated cell populations (Fig. 6A)(Fig. 5E).

The stark differences in the ciliated cell population’s response to SARS-CoV-2 prompted us to compare the ciliated cell types to understand the pathogenic phenotype of the FOXJ1^low^ ciliated cells. We found that FOXJ1^low^ ciliated cells were more metabolically active, while FOXJ1^high^ ciliated cells were more active in cilium motility and ciliogenesis. Other studies have shown that SARS-CoV-2 may induce a marginal upregulation in metabolic activity (52, 53). Interestingly, the differences in axoneme assembly, ciliogenesis, dynein binding, and microtubule motor activity mirrored the microanatomy of 9+2 vs. 9+0 cilium modalities (Fig. 6A)(Fig. 6B). FOXJ1^high^ ciliated cells were more similar to the microanatomy of motile cilia. In contrast, FOXJ1^low^ ciliated cells aligned more with primary or nodal cilium (Fig. 6B). Using scanning electron microscopy, other groups have found that low FOXJ1 expression in ciliated cells is associated with a reduction in motile cilia and mucociliary clearance functionality through impairments of cilia sweeping coordination (54, 55). However, our trajectory and cell phase analysis show similar proportions of FOXJ1^low^ cells in mock and infected samples. Previous studies have shown that lung injury, through various means, can cause the dedifferentiation of ciliated cells and reduced FOXJ1 expression (41, 42, 44, 55–57). The process for obtaining primary lung epithelial cells on an air-liquid interface is strenuous in itself and could be a latent factor in these models. This may be important in that FOXJ1^low^ ciliated cells could be transient and only present after lung injury (or cellular stress, in our *in vitro* model); whether or not these ciliated cell types are abundant in healthy lung tissue remains to be seen (15, 17, 18, 58, 59). However, there has been some evidence that ciliated cells with low expression of FOXJ1 are associated with Bronchiectasis, a condition where the walls of the bronchi are thickened from inflammation and infection (60).

We found that traditional gene enrichment methods using a one-sided Fisher’s Exact Test or a Hypergeometric test using differentially expressed genes yielded few enriched pathways. However, updated rank-based methods significantly reduced false negatives due to intercellular and inter-sample variability. In addition, these methods captured significant functional differences in our comparative analyses on the effects of age and infection status (Fig. 7D)(Fig. 7E). This was particularly important when investigating common antiviral signaling pathways.

Interferon signaling through interferon expression and the induction of hundreds of interferon-stimulated genes (ISGs) is well-documented to be an antiviral protective mechanism; ISGs suppress viral replication and activation of downstream immune signaling (6, 11, 39, 40, 61–68). However, many studies have found very few known interferon-related genes to be expressed after SARS-CoV-2 infection (2–6, 11, 68–70). In our study, we utilized a geneset of previously published ISGs with rank-based scoring to examine indirect type I and type III interferon signaling (39, 40). Using this method, we found significant interferon signaling in a broader sense when examining the breadth of genes associated with an interferon response, rather than induction of type I and III interferon genes in isolation (Fig. 7F). After infection, the host response patterns for ciliated cell types were vastly different. FOXJ1^high^ ciliated cells exhibited expression patterns more similar to club cells while FOXJ1^low^ ciliated cells were more similar to basal cells. This suggests that more transient cell types of the lung epithelium may have a muted response to SARS-CoV-2.

Some limitations of this study include limited donors. Thus, while many of the age-related differences are significant for our cohort, larger future sample sizes may give us more insight into these differences. Additionally, this study relies entirely on RNA and does not include any protein data. Without protein data, the actual functional outcomes of SARS-CoV-2 infection cannot be inferred because the presence of RNA does not necessarily correlate with protein expression; SARS-CoV-2 and other viral pathogens typically have mechanisms that interfere with protein translation (71, 72). Nevertheless, our findings provide important insights into the cellular and molecular mechanisms of SARS-CoV-2 infection in lower lung epithelial cells. Specifically, we found that cellular responses to the virus vastly differ between cell types, highlighting the importance of targeted approaches in developing effective therapeutics and prevention strategies for COVID-19.

## Supporting information

SFig1

SFig2

SFig3

SFig4

SFig5

SFig6

SFig7

SFig8

SFig9

SFig10

## Data availability statement

The datasets presented in this study were deposited in the National Center for Biotechnology Information’s Gene Expression Omnibus database and are available upon request.

## Acknowledgments

The authors want to acknowledge the contributions of Gloria Pryhuber and the LungMAP Biorepository (BRINDL), John Ashton and Michelle Zanche from the UR Genomics Research Center (GRC), Martin Pavelka Jr. and Sonia Rosenberger from the UR Biosafety Level 3 (BSL3) facility and the UR’s Institutional Biosafety Committee (IBC). In addition, the authors would like to acknowledge the advice and support of Mukta G. Palshikar.

## Supplementary Figures

**SFig. 1. Cell cluster distribution.** (A) UMAP of cells colored by sample. (B and C) UMAPs of cells are colored by cluster and split by treatment status (B) and age category (C).

**SFig. 2. Cell cycle phase scoring.** (A) UMAP of cells colored by cell cycle phase. (B) Stacked bar plot of cell cycle proportions by sample. (C) Stacked bar plot of cell cycle proportions by cluster. (D and F) UMAPs of cells are colored by cell cycle phase and split by age category (D) and treatment status (F). (E and G) Stacked bar plots of cells colored by cell cycle phase and divided by age category (E) and treatment status (G). (H) Heatmap of integrated gene expression of highly variable cell cycle markers, annotated by cluster and cell cycle phase assignment.

**SFig. 3. Lung epithelial cell pseudotime comparisons.** (A and B) Boxplots of the range of pseudotime in each cluster, split by age category (A) and treatment status (B).

**SFig. 4. Cell-cell communication frequency.** (A-D) Heatmap of the frequencies of interactions for each pair of potentially communicating cell types. Annotation bar plot on top (“receiving”) and right (“sending”) is the total number of interactions per cell type subsetted by mock samples (A), SARS-CoV-2-infected samples (B), pediatric samples (C), and adult samples (D).

**SFig. 5. Overview of context factorization.** “Age” and “Treatment” bar plots refer to the related context loadings. The “Interactions” histogram is the frequency of individual ligand-receptor pairs. “Senders” is the bar plot of loadings from sender cells. “Receivers” is the bar plot of loadings from receiving cells.

**SFig. 6. Age-related comparative context factorization.** Boxplots of treatment (A) and age-related (B) factor loadings.

**SFig. 7. Heatmaps of factor loadings.** (A) Sample-related factor loadings. (B and C) Sender and receiver loadings for factors 2 (B) and 10 (C).

**SFig. 8. Age-related analysis of combined data.** (A -C) Upset plot of differentially expressed genes in pediatric vs adult donors present in each cluster subsetted by mock-infected samples (A), SARS-CoV-2-infected samples (B), and all samples combined (C). (D) Heatmap of log10 transformed adjusted p-values of enriched KEGG pathways from differentially expressed genes in pediatric vs. adult samples. (E) Dot plot of gene ontology biological process GO terms differentially expressed genes in pediatric vs adult donors present in each cluster.

**SFig. 9. SARS-CoV-2 entry factor expression.** (A) Dotplot of the percentage of cells expressing the indicated SARS-CoV-2-related host entry factors split by mock-infected samples (CTRL, green) and SARS-CoV-2-infected samples (purple). (B) UMAP of log2 normalized gene expression of indicated SARS-CoV-2-related host entry factors.

**SFig. 10. Exploring the pathogenic phenotype of ciliated cells.** (A and B) UMAPs (A) and violin plots (B) of log2 normalized expression of canonical ciliated cell markers.

