## Supplementary figures and images for "Single-cell analysis of lung epithelial cells reveals age and cell population-specific responses to SARS-CoV-2 infection in ciliated cells"

### SFig1

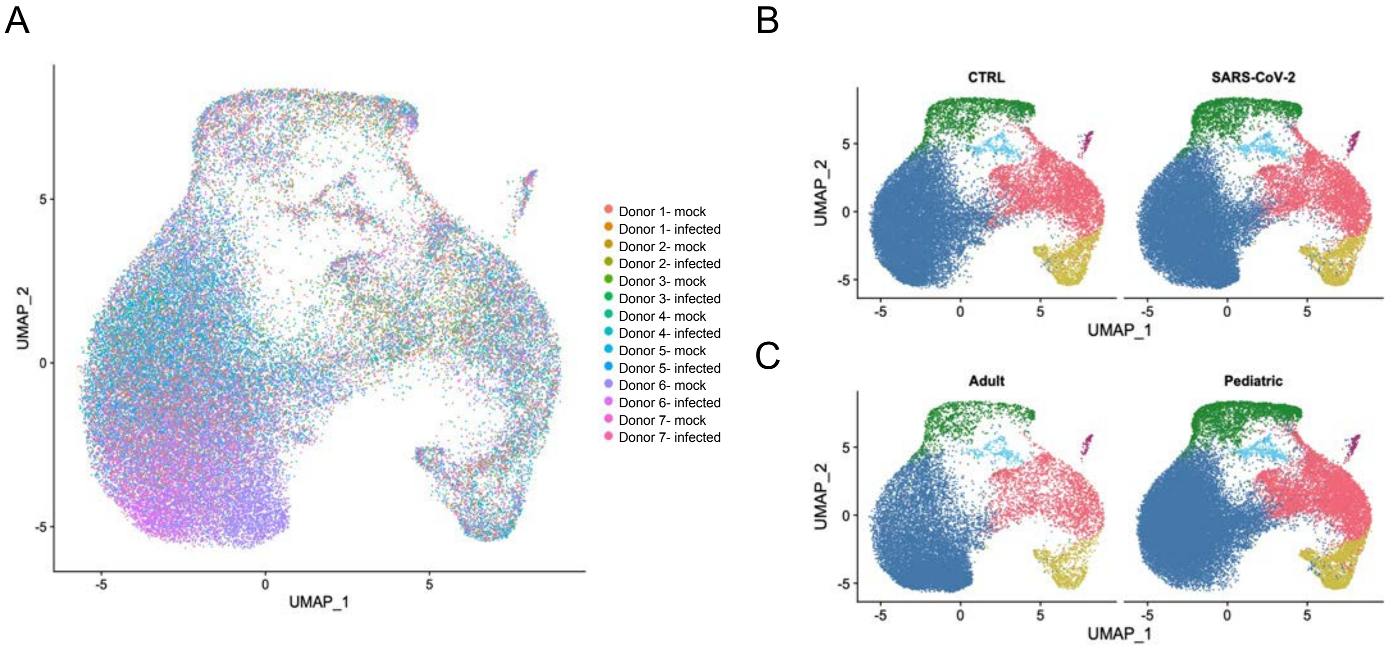

### SFig2

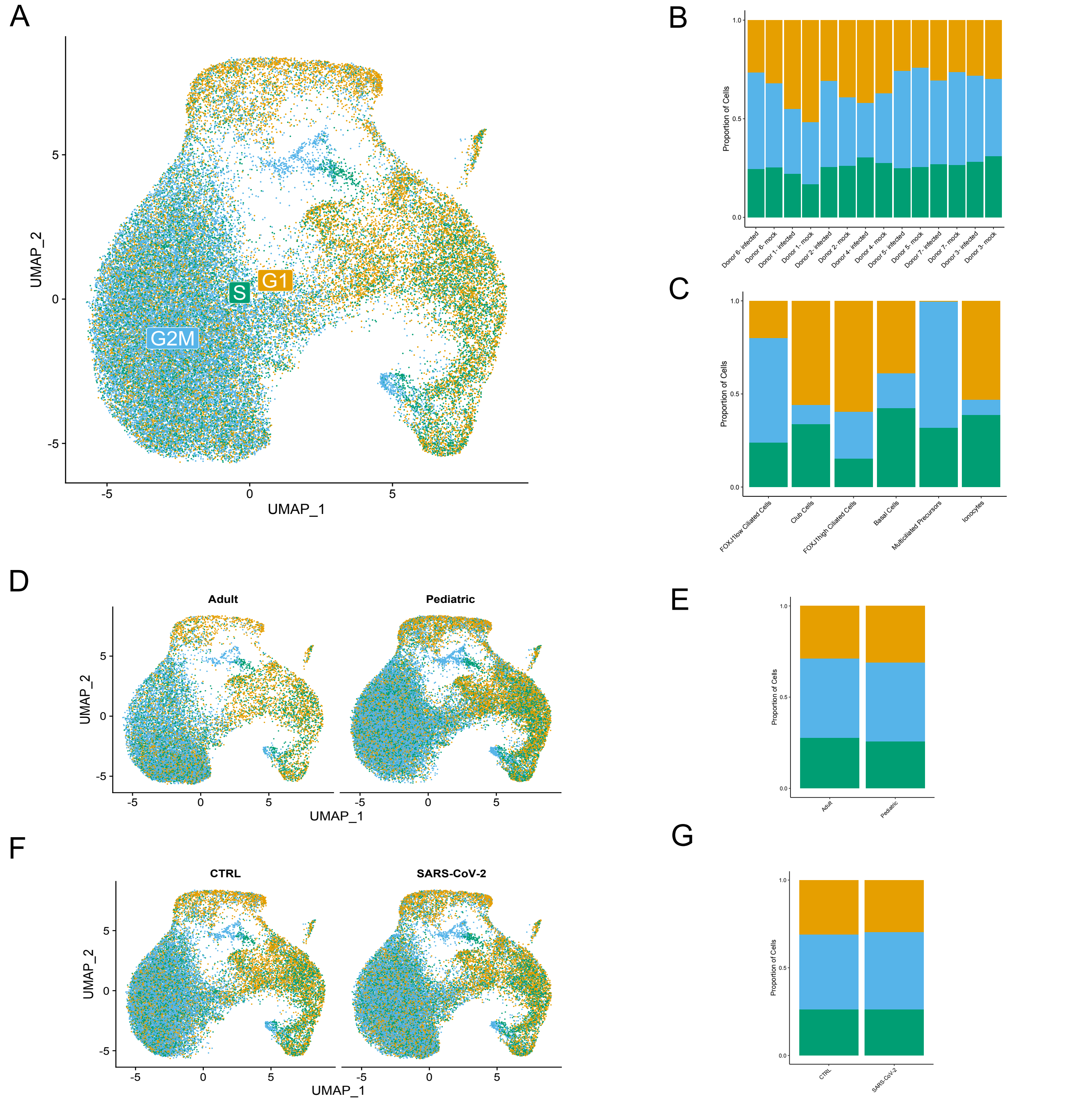

### SFig3

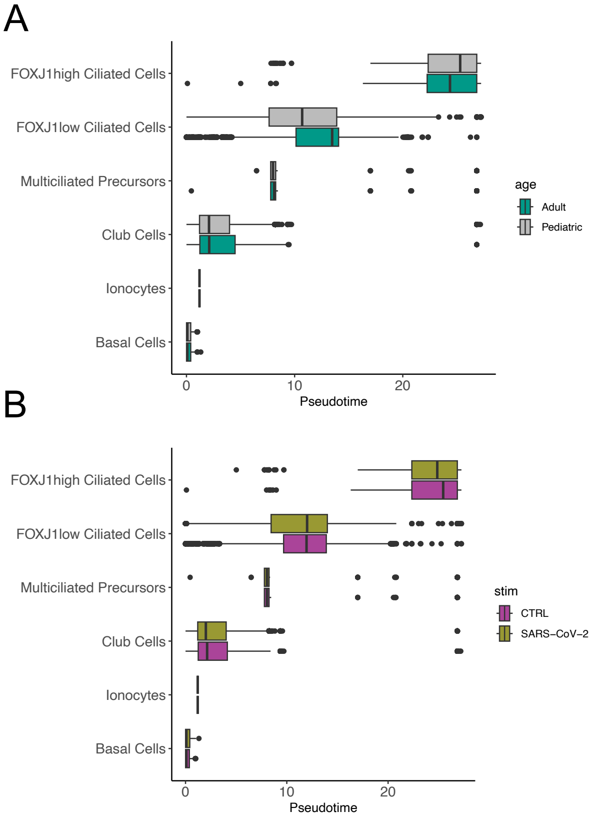

### SFig4

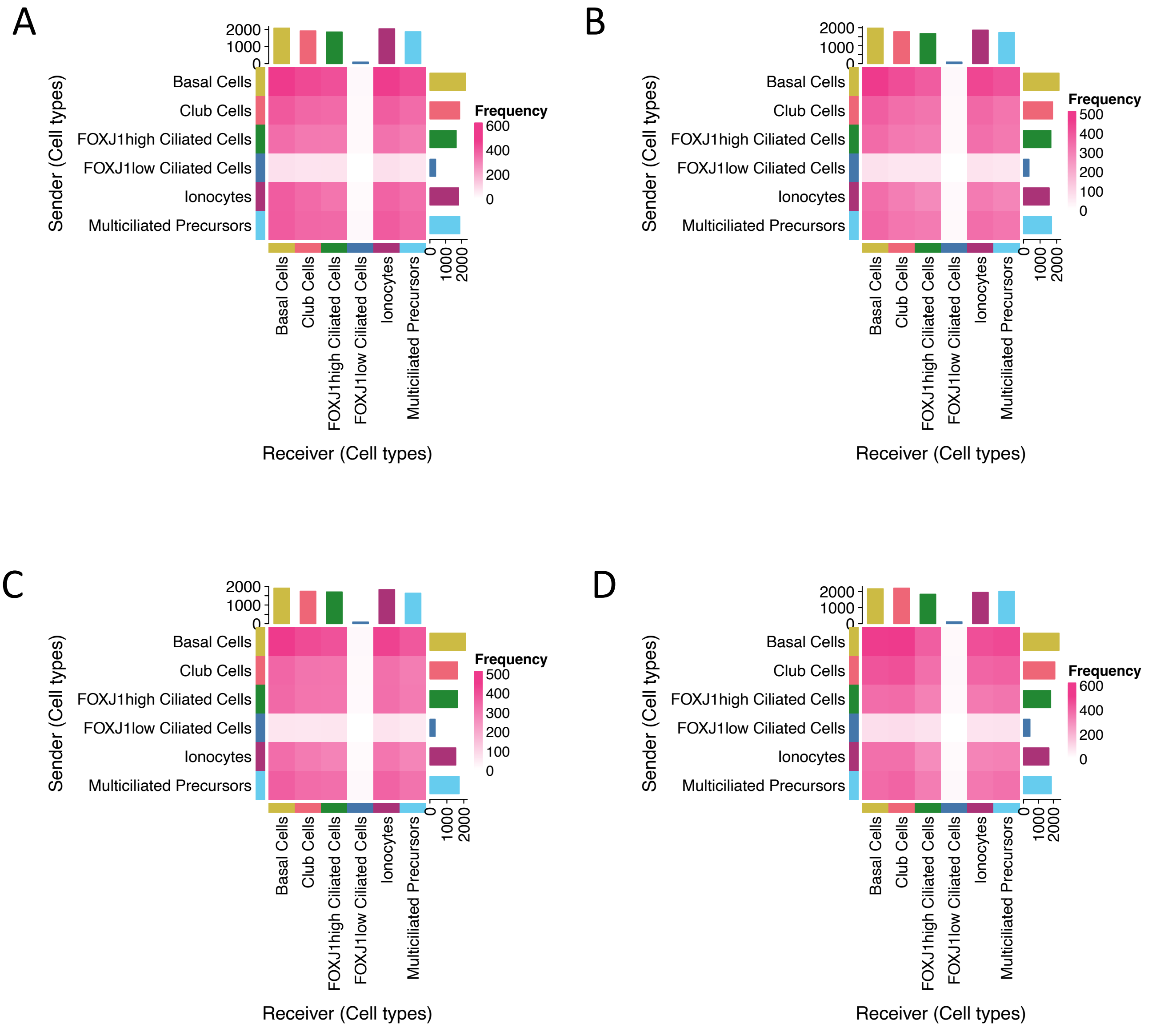

### SFig5

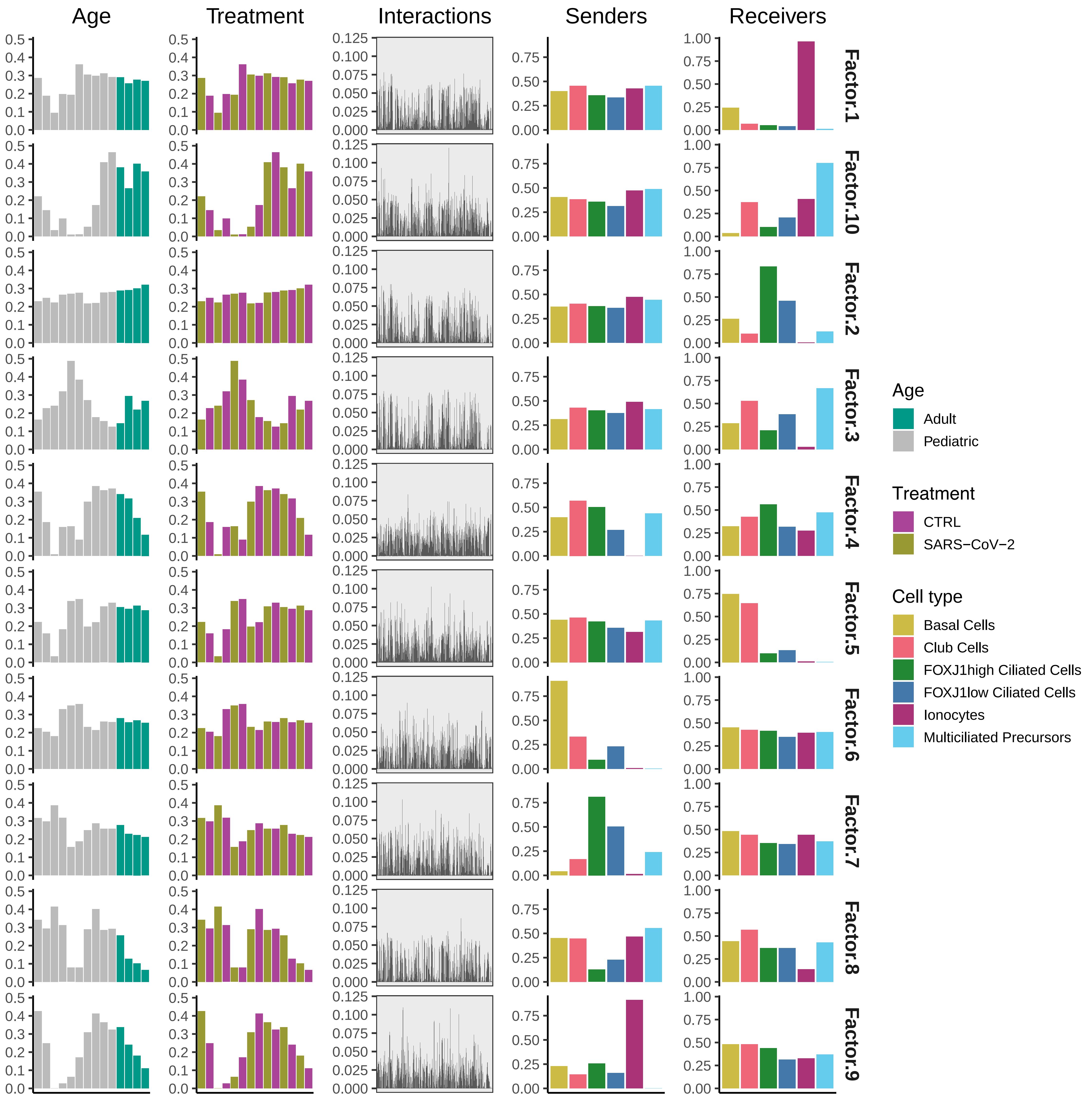

### SFig6

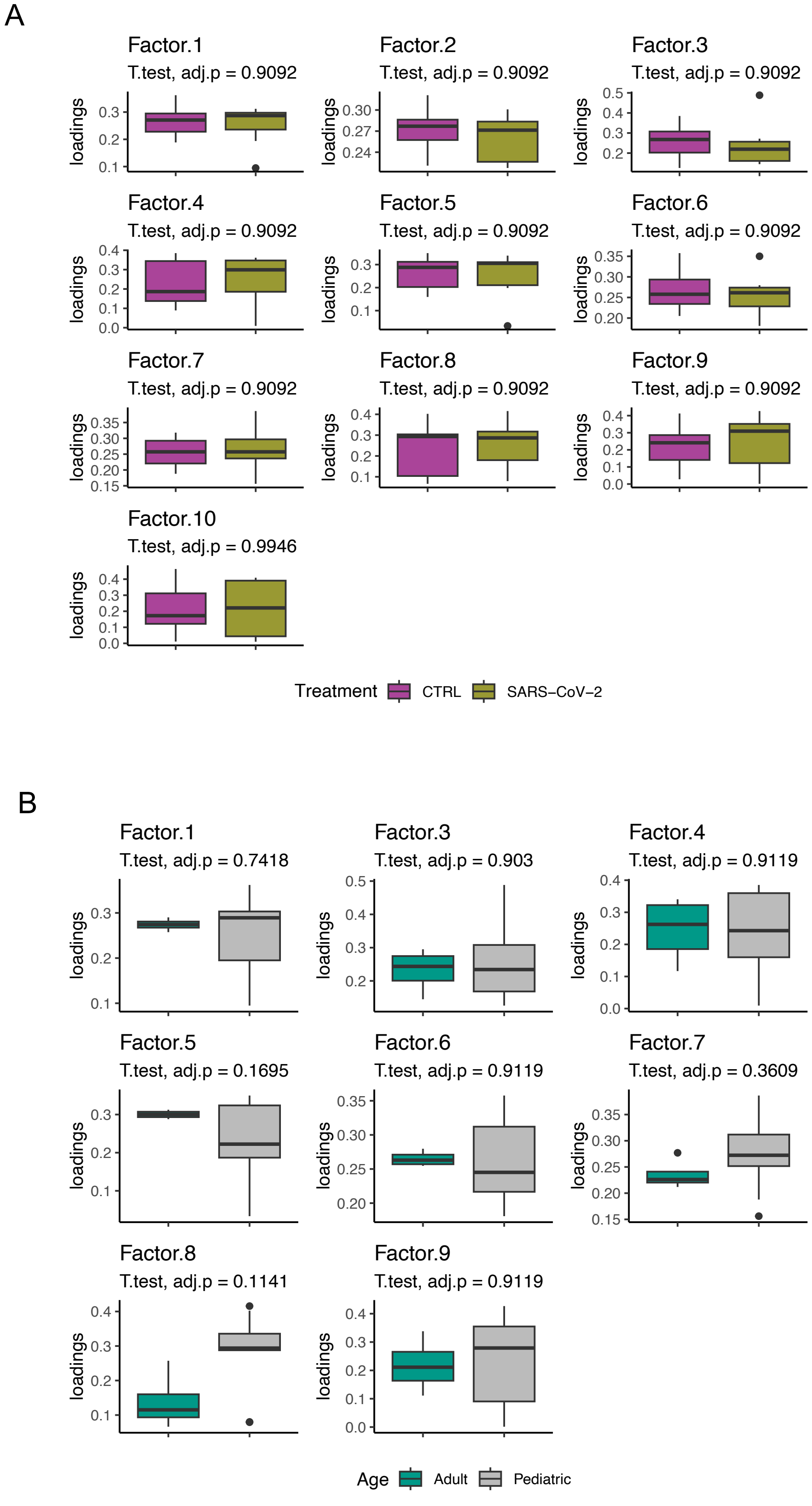

### SFig7

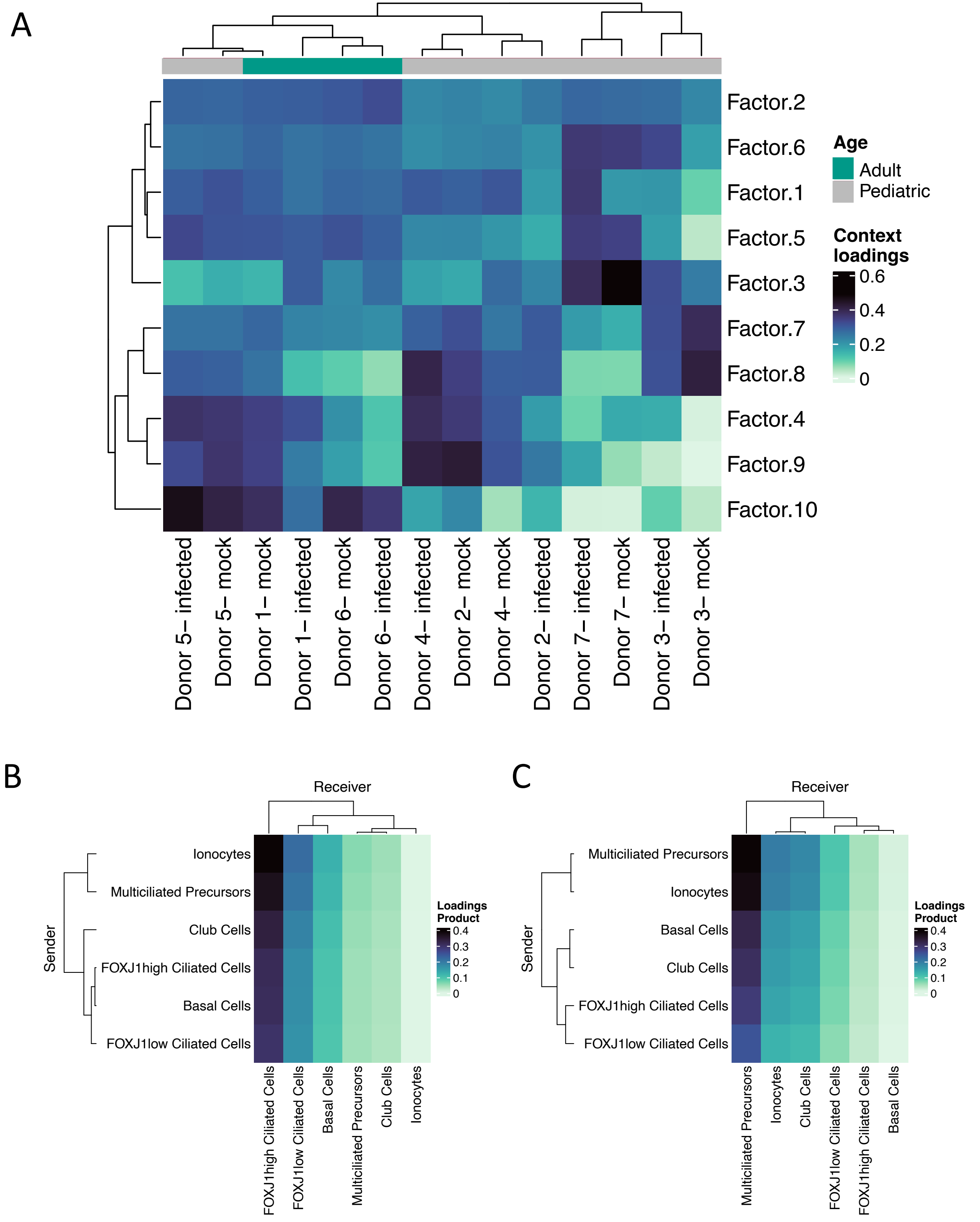

### SFig8

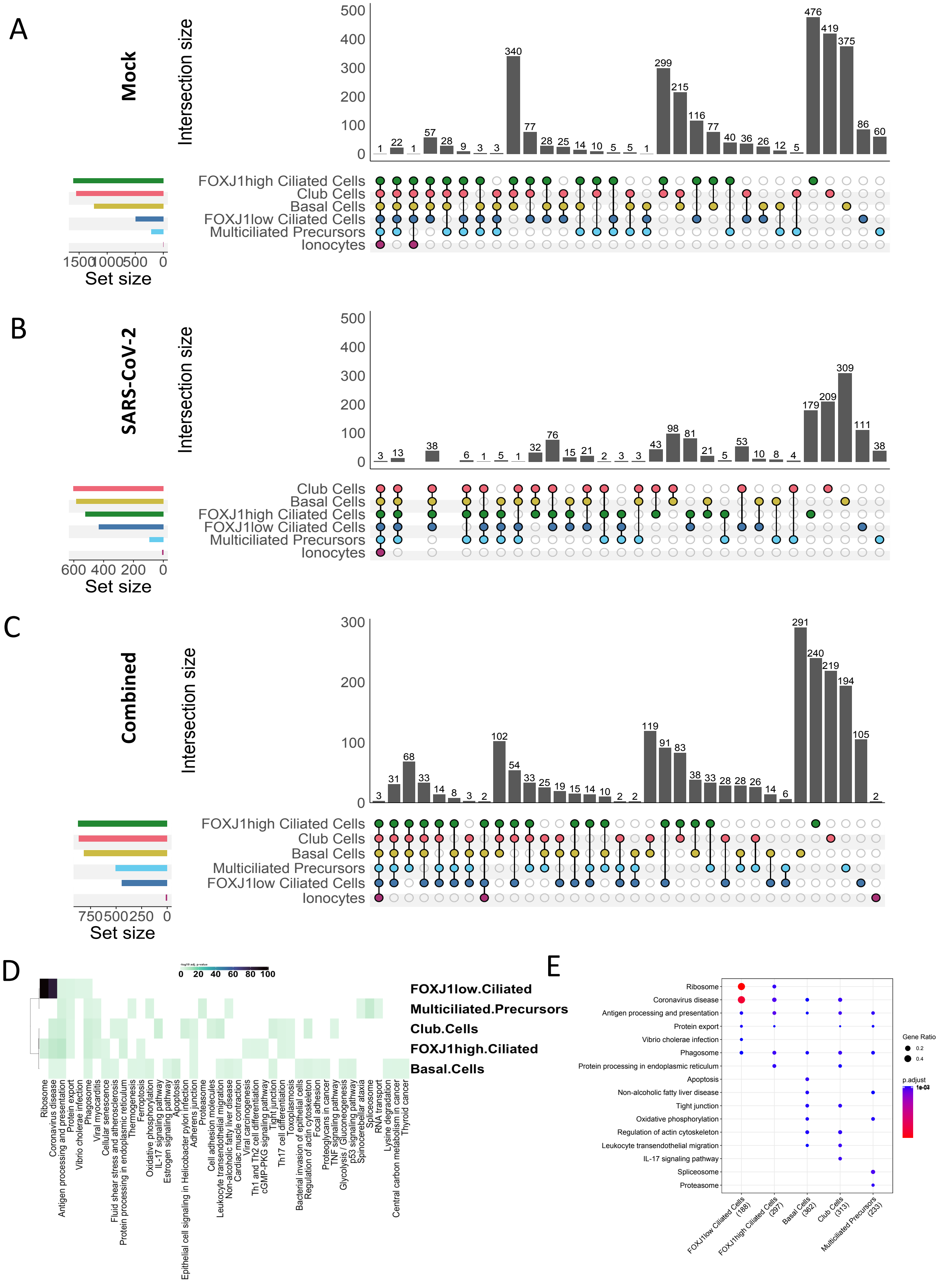

### SFig9

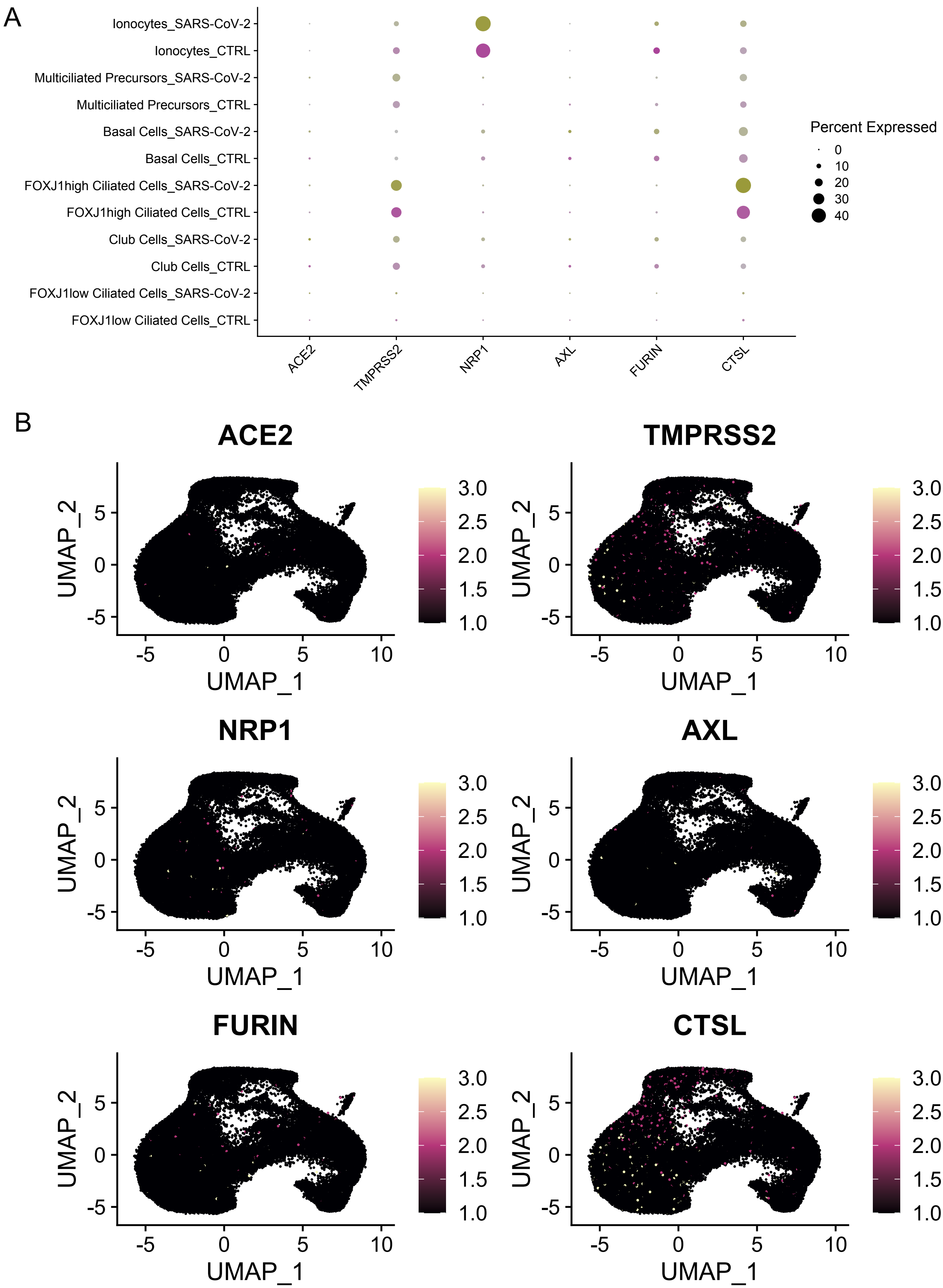

### SFig10

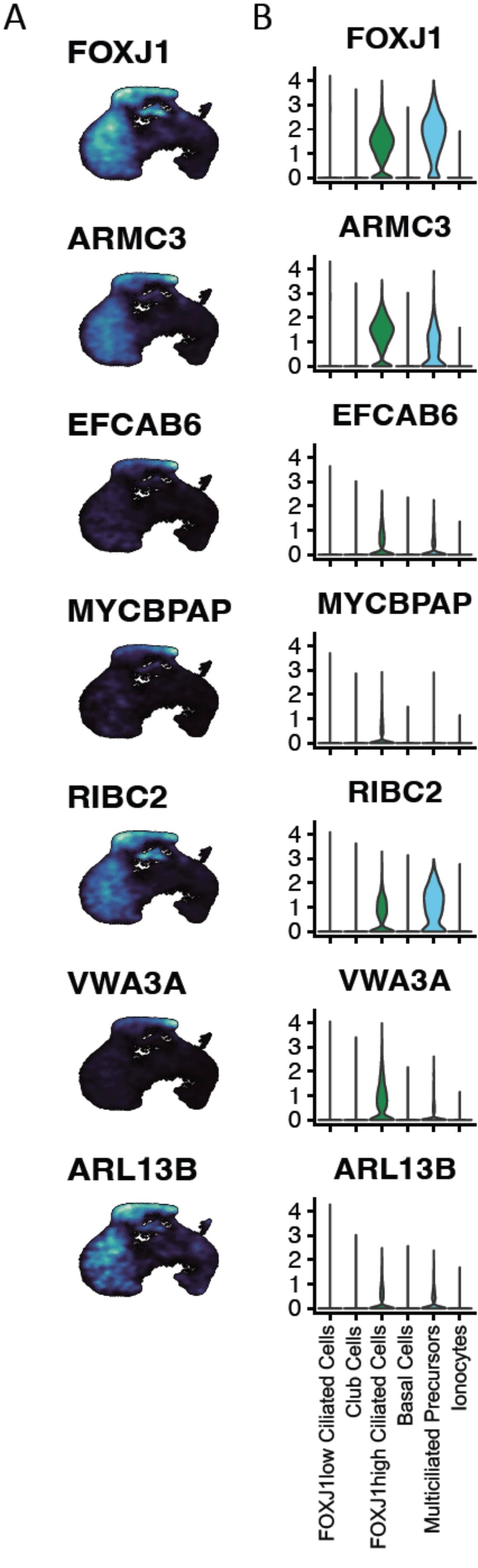
